# Glucagon-like peptide-1 receptor in the human hypothalamus is associated with body mass index and colocalizes with the anorexigenic neuropeptide nucleobindin-2/nesfatin-1

**DOI:** 10.1101/2022.10.22.513332

**Authors:** Aristea Psilopanagioti, Sofia Nikou, Souzana Logotheti, Marina Arbi, Dionysios V. Chartoumpekis, Helen Papadaki

## Abstract

**Introduction:** Glucagon-like peptide-1 (GLP-1) anorexigenic and anti-obesogenic effects are centrally mediated. Data on animals emphasize the importance of neuronal GLP-1 receptor (GLP-1R) for feeding suppression, although it is unclear whether astrocytes participate in the transduction of anorectic GLP-1R-dependent signals. In humans, the brain circuitry underlying these effects remains insufficiently investigated. GLP-1R neuroanatomic localization in human hypothalamus, a brain region with a pivotal role in energy homeostasis regulation, is essential in order to improve our understanding of GLP-1 signaling pathways and central metabolic functions. The present study aimed to explore GLP-1R protein expression in human hypothalamus and its correlation with body mass index (BMI).

**Methods:** Sections of hypothalamus from 28 autopsy cases, 11 with normal weight (BMI < 25 Kg/m^2^) and 17 with non-normal weight (BMI ≥ 25 Kg/m^2^), were examined using immunohistochemistry and double immunofluorescence labeling.

**Results:** Prominent GLP-1R immunoexpression was detected in neurons of several hypothalamic nuclei, including paraventricular, supraoptic, and infundibular nuclei, lateral hypothalamic area (LH), and basal forebrain nuclei. Interestingly, in LH, GLP-1R protein expression was significantly decreased in individuals with BMI ≥ 25 Kg/m2, compared with normal weight counterparts (p=0.03). Furthermore, GLP-1R was moderately and negatively correlated (τb=-0.347, p=0.024) with BMI levels only in the LH. GLP-1R extensively colocalized with the anorexigenic and anti-obesogenic neuropeptide nucleobindin-2/nesfatin-1, but not with the astrocytic marker glial fibrillary acidic protein (GFAP).

**Conclusion:** These data suggest a potential role for GLP-1R in the regulation of energy balance in human hypothalamus, possibly through interactions with nesfatin-1. In LH, an appetite- and reward-related brain region, reduced GLP-1R immunoexpression may contribute to dysregulation of homeostatic and/or hedonic feeding behavior. GLP-1R colocalization with nesfatin-1 in the basal forebrain, a cognition-related brain area, might give impetus towards elucidating additional central actions of GLP-1R.

## Introduction

The incretin hormone glucagon-like peptide-1 (GLP-1) is secreted from intestinal enteroendocrine L cells in response to nutrient ingestion and potentiates insulin release, suppresses glucagon secretion, delays gastric emptying, increases satiety, and reduces appetite and body weight [1]. GLP-1 anorexigenic effect is transduced via GLP-1 receptor (GLP-1R) signaling in several brain areas implicated in the control of energy homeostasis, including the hypothalamus [2]. Although a considerable number of studies have investigated the peripheral actions of GLP-1 in humans [2], information about the effects of GLP-1 and the expression of GLP-1R in the human central nervous system is limited. GLP-1R has been localized in human hypothalamus, using *in situ* hybridization techniques [3, 4] and immunohistochemistry [5]. In the aforementioned studies, GLP-1R showed a particularly heterogenous mRNA and protein expression with strong interindividual variations. Interestingly, a decreased expression of GLP-1R in the paraventricular (PVN) and infundibular (IFN) (arcuate) hypothalamic nuclei was observed in patients with type 2 diabetes mellitus [4]. Furthermore, according to Ten Kulve et al. [4], GLP-1R was sporadically colocalized with energy balance-related neuropeptides, such as neuropeptide Y (NPY), agouti-related peptide (AgRP) and proopiomelanocortin (POMC), in the human IFN.

The astrocytes, that represent an important central nervous system cell population, have been proposed to contribute to the transduction of anorectic GLP-1R-dependent signals in rats [6]. GLP-1R expression in astrocytes has been described within rat brain [6–8] and in a human astrocyte cell line [9]. On the other hand, no colocalization of GLP-1R with the astrocytic marker glial fibrillary acidic protein (GFAP) was observed in any area of the mouse brain examined, including the hypothalamus [10].

In humans, GLP-1R activation reduces neuronal responses to food cues in appetite- and reward-related brain areas, resulting in feeding suppression [11]. Furthermore, it has been suggested that vagal afferents are involved in mediating the effects of GLP-1 on food intake [12]. However, it is unknown whether central GLP-1R acts directly and/or recruits other neuropeptide mediators to exert its anorexigenic and anti-obesogenic effects in humans [2]. Interestingly, nesfatin-1, an 82-amino acid peptide, derived from the precursor nucleobindin-2 (NUCB2), is highly expressed both in human and rodent hypothalamus and exerts pleiotropic actions including an inhibitory effect on food intake and on body weight gain [13–16]. Nesfatin-1 neurons have been shown to contribute to the anorexigenic effects of GLP-1 in an animal model of GLP-1R-mediated feeding suppression [17, 18]. In rats, intraperitoneal administration of GLP-1, resulted in a significant increase of the number of activated nesfatin-1 neurons in hypothalamic nuclei and GLP-1-induced food intake reduction was significantly attenuated by pretreatment with intracerebroventricular administration of antisense nesfatin-1 [19].

Given the emerging evidence that brain circuits regulate energy homeostasis and body weight and the increasing use of GLP-1-based therapies [2], further studying of GLP-1R expression variations in key cell subpopulations of the human hypothalamus is of clinical interest. The present study investigated the association between GLP-1R immunoexpression and clinicopathological data, including body weight. In addition, we examined the colocalization of GLP-1R with the potent anorexigenic neuropeptide NUCB2/nesfatin-1 as well as with the astrocytic marker GFAP.

## Materials and Methods

### Tissue Collection

Hypothalami, from the level of optic chiasm to mammillary bodies, were dissected at autopsy, fixed in neutrally buffered 10% formalin, dehydrated, and paraffin-embedded. Twenty-eight subjects (20 males and 8 females, aged 22-86 years) with no history or postmortem evidence of neuropsychiatric disease were analyzed. Exclusion criteria included: age less than 18 years old, history of neurological and psychiatric disorders, diabetes mellitus, postmortem delay of more than 24 hours, trauma to the brain, use of neuroleptic and anti-obesity medication prior to death, and exposure to toxins. Cases were classified into two groups, according to body mass index (BMI), as follows: “normal weight”, including subjects with BMI < 25 kg/m^2^, and “non-normal weight”, with subjects having a BMI ≥ 25 kg/m^2^ (being overweight or obese). BMI of the examined cases varied between 21.5 and 46.3 kg/m^2^. Post-mortem interval ranged from 3 to 24 hours. Clinicopathological data of examined cases are presented in Table 1. The study protocol was reviewed and approved by the Ethics Committee of the University Hospital of Patras (approval number 94/10.5.07). The research was conducted ethically in accordance with the World Medical Association Declaration of Helsinki.

**Table 1:** Clinicopathological data.Abbreviations: BMI, Body Mass Index; PMI, Post Mortem Interval.

| Case | Sex | Age (years) | BMI (Kg/m<sup>2</sup>) | PMI (hours) | Cause of death, Medical History |
| --- | --- | --- | --- | --- | --- |
| 1 | M | 30 | 21.5 | 10 | Homicide |
| 2 | F | 65 | 21.6 | 24 | Automobile accident |
| 3 | M | 66 | 23.0 | 16 | Cardiac tamponade, myocardial infarction, hypertension |
| 4 | M | 53 | 23.0 | 10 | Myocardial infarction |
| 5 | M | 40 | 23.3 | 24 | Myocardial infarction |
| 6 | M | 57 | 23.9 | 12 | Myocardial infarction |
| 7 | F | 27 | 24.2 | 11 | Automobile accident |
| 8 | M | 31 | 24.8 | 24 | Dilated cardiomyopathy |
| 9 | F | 55 | 24.9 | 24 | Myocardial infarction |
| 10 | F | 56 | 24.9 | 11 | Myocardial infarction |
| 11 | M | 24 | 24.9 | 5 | Automobile accident |
| 12 | M | 64 | 27.7 | 17 | Myocardial infarction |
| 13 | M | 37 | 27.8 | 3 | Automobile accident, hypertension |
| 14 | F | 58 | 29.0 | 24 | Automobile accident |
| 15 | F | 77 | 29.1 | 12 | Automobile accident, hypertension |
| 16 | M | 86 | 29.1 | 15 | Gastric cancer/hemorrhage |
| 17 | F | 22 | 29.3 | 8 | Automobile accident |
| 18 | M | 60 | 29.4 | 24 | Automobile accident |
| 19 | F | 64 | 29.4 | 3 | Cor pulmonale |
| 20 | M | 58 | 30.5 | 24 | Myocardial infarction |
| 21 | M | 63 | 30.7 | 12 | Gastric hemorrhage, alcoholism |
| 22 | M | 56 | 31.6 | 12 | Aspiration pneumonia, hypertension |
| 23 | M | 39 | 31.7 | 2 | Myocardial infarction |
| 24 | M | 59 | 32.4 | 12 | Myocardial infarction |
| 25 | M | 27 | 32.4 | 24 | Myocardial infarction, familial hypercholesterolemia |
| 26 | M | 42 | 33.7 | 6 | Myocardial infarction |
| 27 | M | 58 | 35.5 | 10 | Electrocution |
| 28 | M | 54 | 46.3 | 12 | Myocardial infarction |

Abbreviations: BMI, Body Mass Index; PMI, Post Mortem Interval.

### Histochemistry

Standard H&E and luxol fast blue-cresyl violet stains were performed on representative sections for morphological and topographic assessment of brain areas.

### Immunohistochemistry

Consecutive 4-μm coronal sections of hypothalamus (preoptic area, anterior hypothalamic area, tuberal region, and mammillary region) were analyzed. Three to six hypothalamic sections per nucleus per case, taken at regular intervals throughout the rostrocaudal axis of each nucleus, were deparaffinized in a series of xylene and ethanol. Immunohistochemistry was performed as previously described [16, 20]. In brief, after blocking endogenous peroxidase activity, by incubating the sections in methanol containing 0.3 % H2O2 for 20 minutes, and performing antigen retrieval in citrate buffer (pH 6), sections were incubated overnight at 4 °C using a rabbit polyclonal anti-GLP-1R antibody (1:50; Origene, Rockville, MD, USA, TA336864). Specificity of GLP-1R antiserum has been previously described in detail [21–23]. Color was developed using the EnVision Flex Kit (Agilent Technologies Inc., Santa Clara, CA, USA), according to the manufacturer’s instructions. Briefly, after incubation with the primary antibody, sections were washed in TBS and incubated with the secondary antibody coupled with activation reagent EnVision Flex horseradish peroxidase (HRP) for 20 minutes. Visualization was achieved by incubating slides 10 minutes with 3,3’-diaminobenzidine (DAB). Slides were counterstained with Harris’ hematoxylin, dehydrated in ascending alcohol row, and permanently mounted. All sections were simultaneously processed in the presence of appropriate positive and negative controls.

Hypothalamic structures were identified with reference to the atlas of the human brain by Mai J et al. [24]. Qualitative analysis of protein expression was performed independently and graded blindly by two researchers (H.P. and A.P.). Images were captured on 3D HISTECH Ltd Hungary Pannoramic DESK Scanner 1.16 and analyzed using Pannoramic Viewer Software Version 1.15.4 C3D HISTECH Ltd, Hungary.

Immunostaining score was calculated in a semiquantitative fashion, using a four-point scale system, as follows: +++ (more than 50% of cells are positive and staining intensity is high); ++ (more than 50% of cells are positive and staining intensity is moderate); + (more than 50% of cells are positive and staining intensity is low); - (no staining).

In order to further validate the results and confirm specificity, an additional rabbit polyclonal anti-GLP-1R antibody was used (1:50; Merck Millipore, Darmstadt, Germany, AB9433-I).

Pancreatic tissue was used as positive control for GLP-1R. Negative controls were prepared by omitting the primary antibody or by substituting GLP-1R antiserum with normal rabbit serum.

### Double immunofluorescence labeling

To investigate potential overlap between: a) GLP-1R and NUCB2/nesfatin-1 and b) GLP-1R and GFAP immunoexpression, we performed double immunofluorescence analysis of human hypothalamus.

Hypothalamic sections from 8 individuals, 4 with normal and 4 with non-normal BMI (cases 2, 4, 6, 9, 14, 19, 23, 25 in Table 1), were double immunostained using the following antisera couples: a) rabbit polyclonal anti-GLP-1R (1:50; Origene) and mouse monoclonal anti-NUCB2/nesfatin-1 antibody (1:50; Merck Millipore, MABS1164), and b) rabbit polyclonal anti-GLP-1R (1:50; Origene) and mouse monoclonal anti-GFAP antibody (1:500; Merck Millipore, MAB360). Specificity of antibodies used for colocalization studies was previously published [16, 21–23, 25].

Hypothalamic sections were deparaffinized, blocked in phosphate-buffered saline containing 10% fetal bovine serum, 3% bovine serum albumin, and 0.1% Tween-20, for 2 h, and incubated overnight with couples of primary antibodies at 4°C. After washing, sections were incubated at room temperature for 1 h with secondary antibodies Alexa Fluor 568 goat anti-mouse IgG and Alexa Fluor 488 goat anti-rabbit IgG (1:500; Invitrogen Life Technologies, Rockford, IL, USA), counterstained with Hoechst 33258 (1:1,500; Sigma-Aldrich Chemie GmbH, Germany), and coverslipped with fluorescent mounting medium (Merck Millipore). Staining specificity was determined by omission of primary antibodies. Slides were analyzed in the Advanced Light Microscopy Facility of Patras Medical School with the use of a Nikon Eclipse TE 2000-U inverted microscope. Images were captured using the INFINITY software package (Lumenera Corporation, Ottawa, Kanada). In hypothalamic and basal forebrain nuclei, the percentage of double-labeled neurons with GLP-1R and NUCB2/nesfatin-1 was counted by examining ten high-power fields per nucleus per section at a magnification of 400× and averaged to obtain a single value. Figures from double immunofluorescence labeling of GLP-1R and GFAP proteins were obtained on a Confocal Leica SP5 microscope (Leica Microsystems, France).

### Statistical analysis

All data were analyzed with IBM SPSS Statistics for Windows version 25.0 (IBM Corp., Armonk, N.Y., USA). Ordinal variables, such as protein expression-immunoreactivity scores, were presented as frequency distributions, with percentages. Quantitative variables were defined as mean value ± standard error of the mean (SEM). Normality was tested both graphically and using the Shapiro-Wilk test. Normal distribution was not met and therefore non-parametric statistical methods were employed. Group differences in the distribution of immunoreactivity scores were evaluated with Mann–Whitney U test. Correlations between GLP-1R immunoexpression and clinicopathological parameters were examined using the Kendall’s Tau-b correlation coefficient. P<0.05 was considered statistically significant.

## Results

### Distribution of GLP-1R immunoreactivity in human hypothalamus is different between normal weight and overweight or obese subjects

GLP-1R immunoreactive (GLP-1R-ir) neurons exhibited diffuse cytoplasmic immunoperoxidase labeling and widespread distribution in several hypothalamic nuclei and areas (Figure 1). The PVN (magnocellular and parvicellular neuronal populations), supraoptic (SON) (dorsolateral, dorsomedial, and ventromedial parts), and IFN nuclei, the lateral hypothalamic area (LH), and the cholinergic basal forebrain nuclei, including the basal nucleus of Meynert and the diagonal band of Broca, showed the most prominent GLP-1R immunoreactivity. In addition, GLP-1R-ir neurons showing moderate, weak, and in a few cases negative immunostaining were detected in the dorsomedial (DMH), ventromedial (VMH), perifornical, and tuberomammillary (TM), suprachiasmatic, and mammillary hypothalamic nuclei as well as in bed nucleus of the stria terminalis. No immunopositivity for GLP-1R was observed in human glial cells. GLP-1R-ir fibers were widely distributed in the hypothalamus. A schematic representation of GLP-1R-ir cell bodies and fibers is depicted in Figure 2.

**Figure 1.**
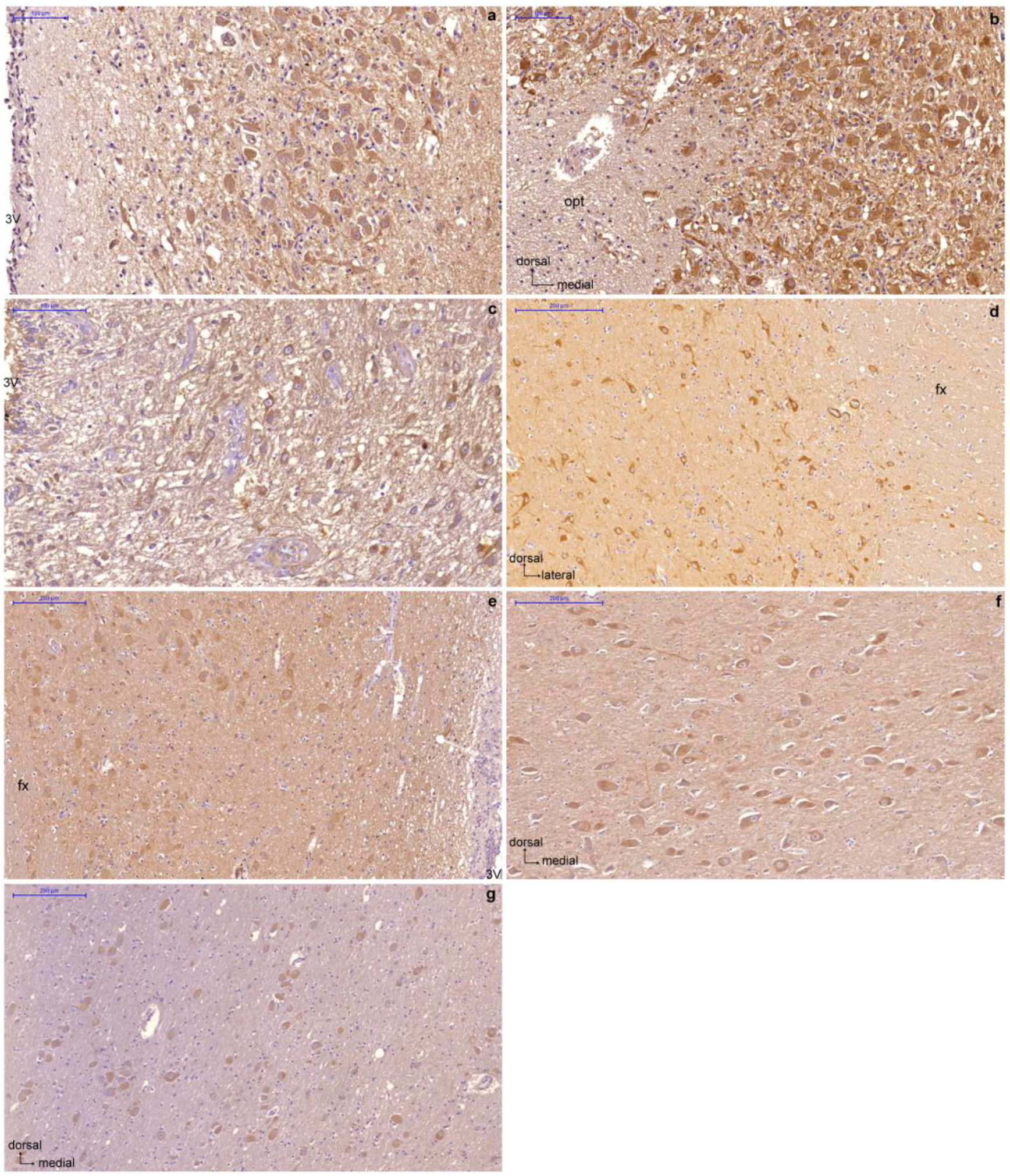
Representative sections of human hypothalamus showing GLP-1R immunoexpression in neurons of the paraventricular nucleus (a), supraoptic nucleus (b), infundibular nucleus (c), lateral hypothalamic area (d), dorsomedial nucleus (e), basal nucleus (f), diagonal band (g). 3V, third ventricle; fx, fornix; opt, optic tract. Scale bars indicate 100 μm (a, b, c) and 200 μm (d, e, f, g).

**Figure 2.**
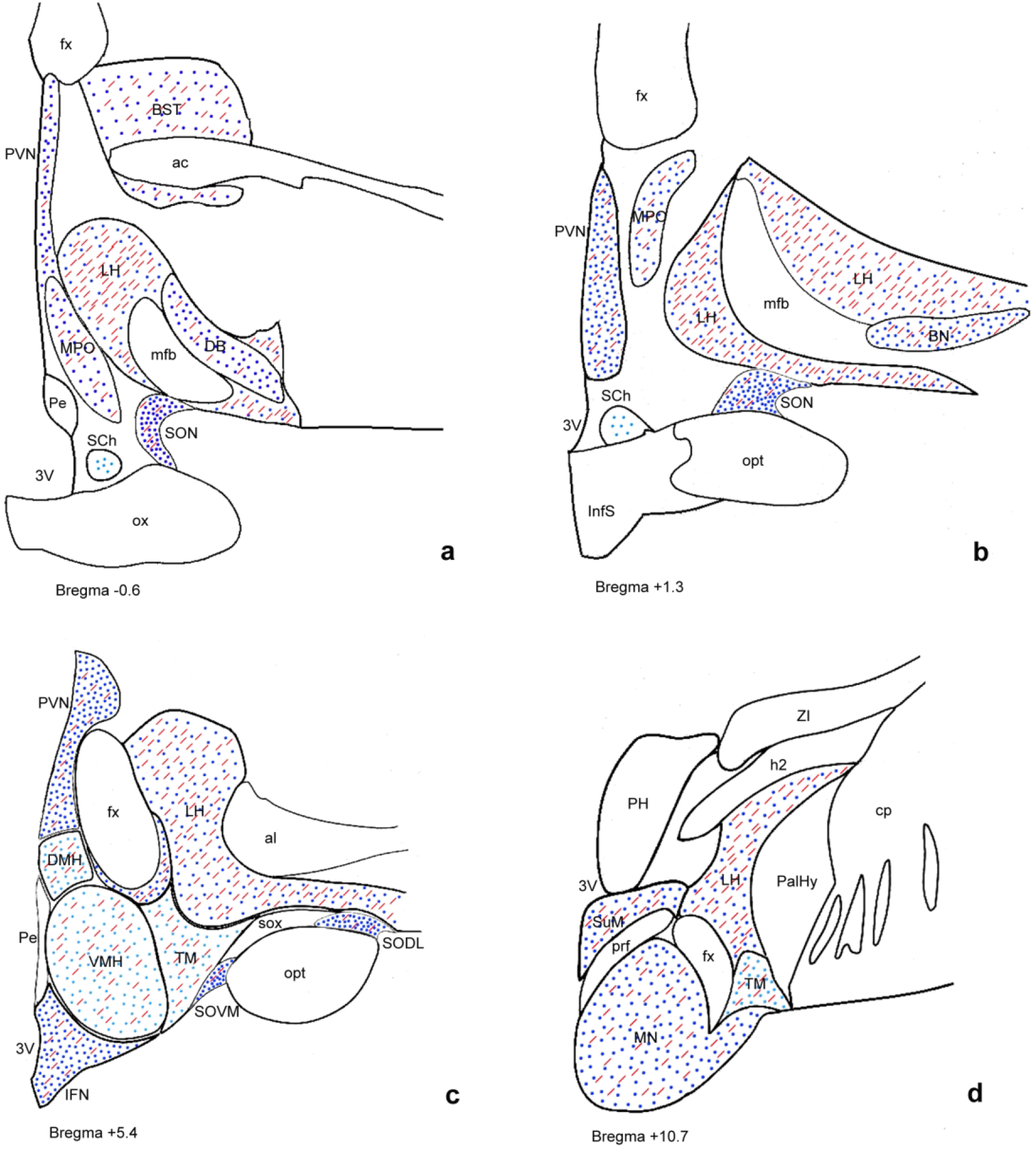
Schematic illustration of GLP-1R distribution in the human hypothalamus. Depiction of GLP-1R-ir cell bodies (blue dots) and GLP-1R-ir fibers (red lines) in four representative coronal sections of human hypothalamus arranged from rostral (a) to caudal (d). Dark blue coloring indicates areas showing high staining intensity for GLP-1R and light blue indicates less intense staining. 3V, third ventricle; ac, anterior commissure; al, ansa lenticularis; BN, basal nucleus; BST, bed nucleus of the stria terminalis; cp, cerebral peduncle; DB, diagonal band; DMH, dorsomedial hypothalamic nucleus; fx, fornix; h2, lenticular fascicle; ic, internal capsule; IFN, infundibular nucleus; InfS, infundibular stalk; LH, lateral hypothalamic area; mfb, medial forebrain bundle; mfb, medial forebrain bundle; MN, mammillary nucleus; MPO, medial preoptic nucleus; opt, optic tract; ox, optic chiasm; PalHy, pallidohypothalamic nucleus; Pe, periventricular nucleus; PeF, perifornical nucleus; PH, posterior hypothalamic area; prf, principal fasciculus; PVN, paraventricular nucleus; SCh, suprachiasmatic nucleus; SON, supraoptic nucleus; SODL, supraoptic nucleus (dorsolateral part); SOVM, supraoptic nucleus (ventromedial part); sox, supraoptic commissure; SuM, supramammillary nucleus; TM, tuberomammillary nucleus; VMH; ventromedial hypothalamic nucleus; ZI, zona incerta.

GLP-1R distribution pattern was highly similar among all subjects, although an interindividual variation in staining intensity was noticed in a few hypothalamic nuclei. Interestingly, in LH, immunohistochemical expression of GLP-1R displayed a significantly different distribution between the two BMI groups. Normal weight subjects showed significantly higher immunoreactivity scores compared to their overweight or obese counterparts with BMI ≥ 25kg/m^2^ (P=0.03). The results of the comparison of immunoreactivity scores between BMI categories are presented in Table 2. Figure 3 illustrates the difference in GLP-1R immunoreactivity score, in LH, between a normal weight (case 4) and an excess adiposity BMI subject (case 28), both males with similar age and post mortem intervals, and the same cause of death. GLP-1R protein expression in all other hypothalamic nuclei was not significantly correlated with BMI weight status category (P>0.05). No statistically significant association between GLP-1R immunoexpression and sex, age, postmortem interval or cause of death was noticed (P>0.05). Regional differences in GLP-1R immunostaining scores for all examined cases are summarized in Supplemental Tables 1 and 2.

**Figure 3.**
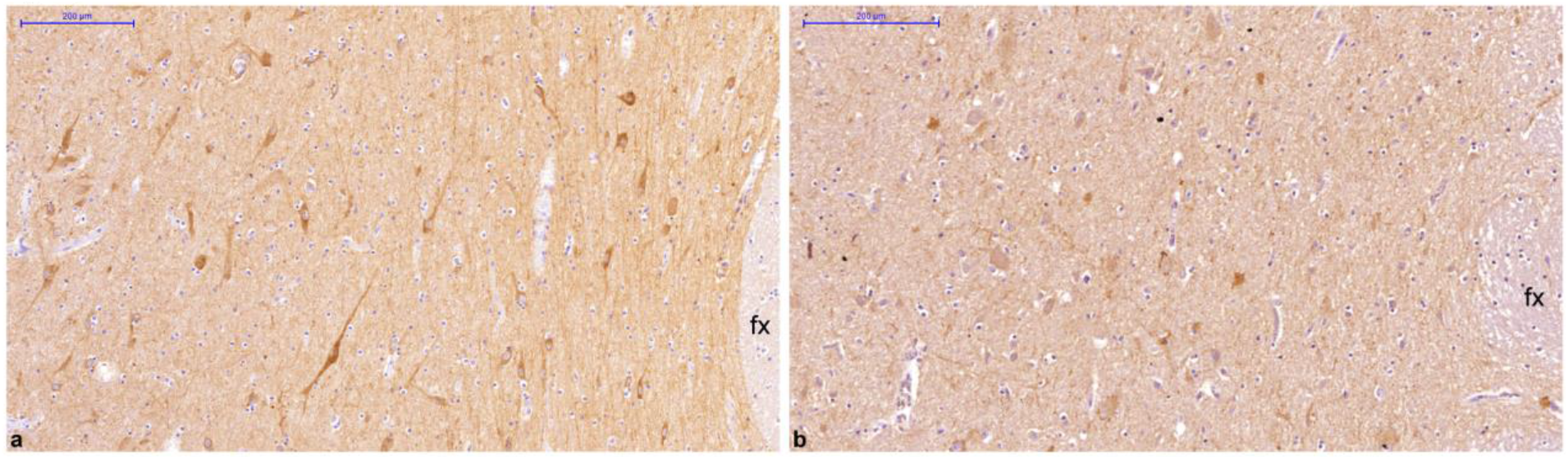
Comparison of GLP-1R immunoexpression in lateral hypothalamic area from a normal-weight and a non-normal weight individual, both with similar clinicopathological characteristics. Images, taken in similar rostrocaudal levels (medial zone of the tuberal region), are derived from a male with BMI 23.0 kg/m^2^, displaying high GLP-1R immunoreactivity (a) and a male with BMI 46.3 kg/m^2^, showing low GLP-1R immunoreactivity (b). Bar denotes 200 μm.

**Table 2:**
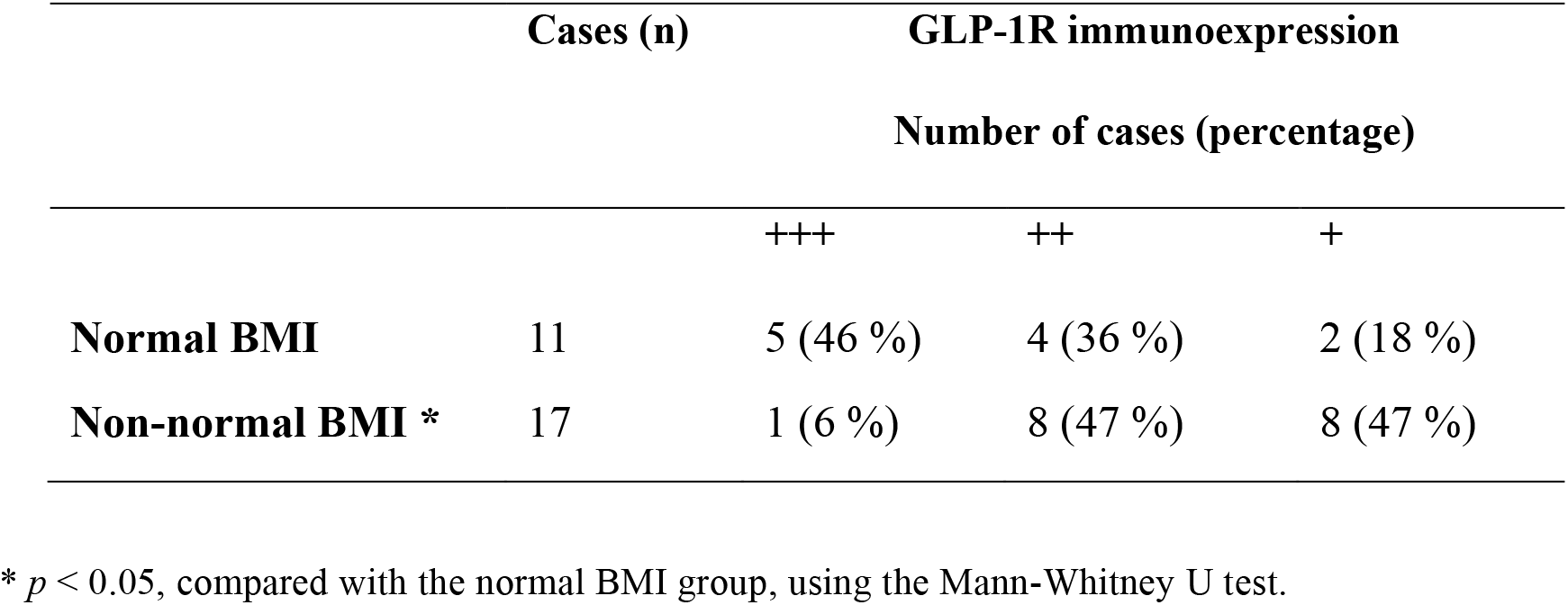
Comparison between the semi-quantitatively expressed GLP-1R immunoreactivity scores of normal body mass index (BMI) (<25 kg/m^2^), and non-normal BMI (≥25 kg/m^2^) cases in the lateral hypothalamic area. Immunoreactivity score is expressed as follows: +++, >50% of cells exhibit strong immunoexpression; ++, >50% of cells exhibit moderate immunoexpression; +, >50% of cells exhibit weak immunoexpression.

Furthermore, GLP-1R immunoexpression did negatively correlate with BMI in LH (Kendall’s Tau-b=-0.347, p=0.024). BMI was not correlated with GLP-1R in any other hypothalamic nuclei/areas analyzed in the present study (P>0.05) (Table 3). In addition, no statistically significant correlation between GLP-1R and clinicopathologic parameters, such as sex, age, cause of death, and postmortem interval, was observed (P>0.05).

**Table 3:**
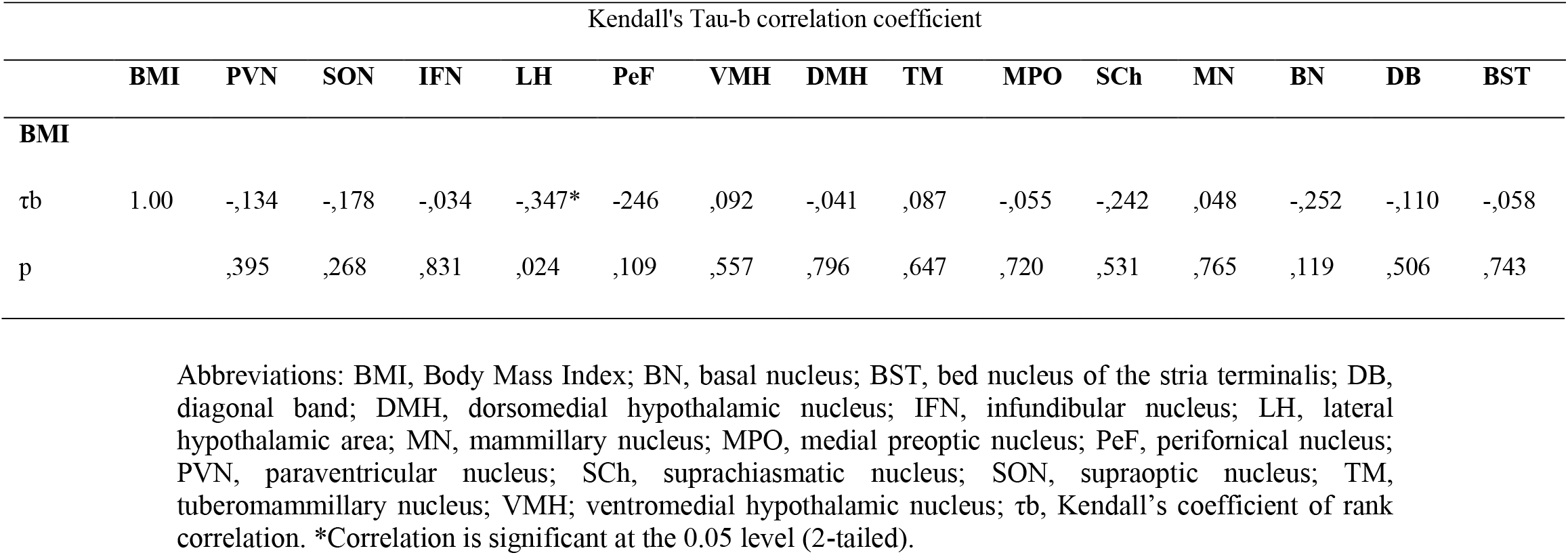
Correlations between BMI and expression of GLP-1R in human hypothalamic nuclei.

Moreover, we verified our results regarding GLP-1R expression by using another antibody against GLP-1R (Merck-Millipore) that targets a non-overlapping epitope instead of the one originally used (Origene). Similar results regarding the GLP-1R topography and staining intensity in hypothalamus were obtained using both antibodies, as shown in Supplemental Table 2.

For control purposes the GLP-1R expression was examined in the pancreas. As expected, intense GLP-1R immunoexpression was observed in human pancreas. Omitting the primary antibody or substituting GLP-1R antiserum with normal rabbit serum resulted in no immunoreactivity in the human pancreas and hypothalamus.

### GLP-1R colocalizes with anorexigenic and anti-obesogenic neuropeptide NUCB2/nesfatin-1, but not with the astrocytic marker glial fibrillary acidic protein (GFAP)

GLP-1R colocalized extensively with NUCB2/nesfatin-1 in several hypothalamic nuclei, namely PVN, SON, IFN, LH, and in basal forebrain nuclei (Figure 4). In the PVN, the vast majority of GLP-1R-ir neurons overlapped with NUCB2/nesfatin-1 (91.2 ± 2.4 %). Numbers represent mean value ± Standard Error of the Mean (SEM). On average, in the SON, of all GLP-1R-ir perikarya 96.7 ± 0.9 % exhibited NUCB2/nesfatin-1 immunoreactivity. Similarly, in the IFN, GLP-1R-ir neurons colocalized with NUCB2/nesfatin-1 at a percentage of 89.7 ± 3.9 %. In LH, of all GLP-1R-ir neurons, 97.0 ± 1.9 % contained NUCB2/nesfatin-1. Furthermore, in basal nucleus, 95.9 ± 1.4 % of GLP-1R-ir cells overlapped with NUCB2/nesfatin-1. Conversely, the percentages of NUCB2/nesfatin-1-ir neurons coexpressing GLP-1R, in the PVN, SON, IFN, LH, and basal nucleus, were 55.1 ± 2.8 %, 60.3 ± 4.3 %, 62.2 ± 4.0 %, 57.7 ± 6.5 %, and 75.3 ± 7.7 %, respectively. Percentages of GLP-1R-ir neurons colocalizing with NUCB2/nesfatin-1 as well as of NUCB2/nesfatin-1-ir neurons colocalizing with GLP-1R are shown in Tables 4 and 5, respectively. In all hypothalamic nuclei/areas examined, comparing medians of percentages of GLP-1R-ir neurons colocalized with NUCB2/nesfatin-1 as well as of NUCB2/nesfatin-1-immunopositive neurons coexpressing GLP-1R revealed no statistically significant difference between normal weight and non-normal weight BMI groups (P> 0.05). Furthermore, GLP-1R did not colocalize with GFAP in any of the examined cases (Figure 5). Omission of the primary antibodies resulted in signal absence.

**Figure 4.**
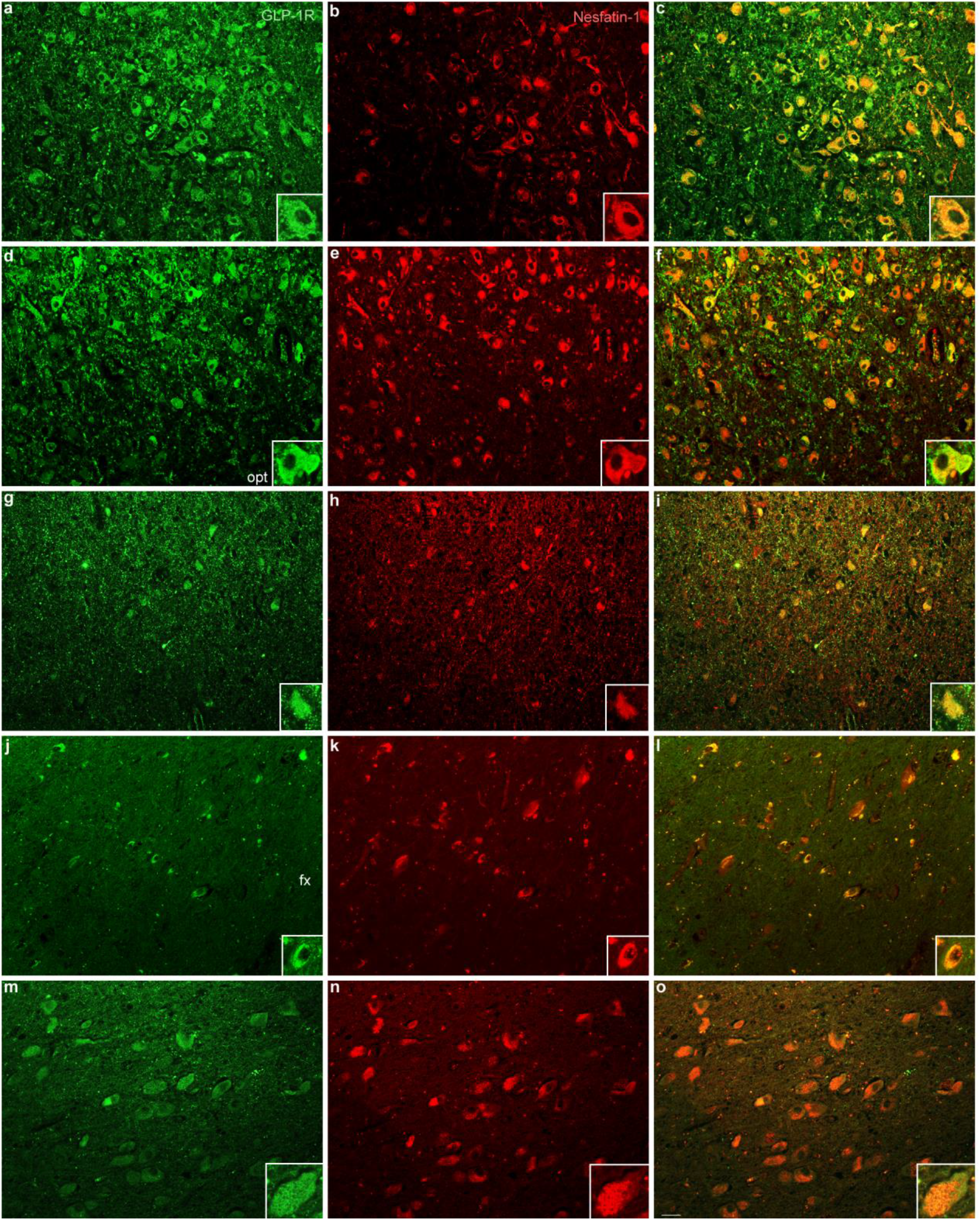
Representative light microscopic fluorescent images depicting GLP-1R and NUCB2/nesfatin-1 colocalization in the human hypothalamus. Representative sections of paraventricular nucleus (a-c), supraoptic nucleus (d-f), infundibular nucleus (g-i), lateral hypothalamic area (g-l), and basal nucleus (m-o) double stained for GLP-1R (green) and NUCB2/nesfatin-1 (red). Merged images depict double stained neurons (c, f, i, l, o) (200x). Insets represent neurons at higher magnification (400x). fx, fornix; opt, optic tract. Scale bar indicates 50 μm.

**Figure 5.**
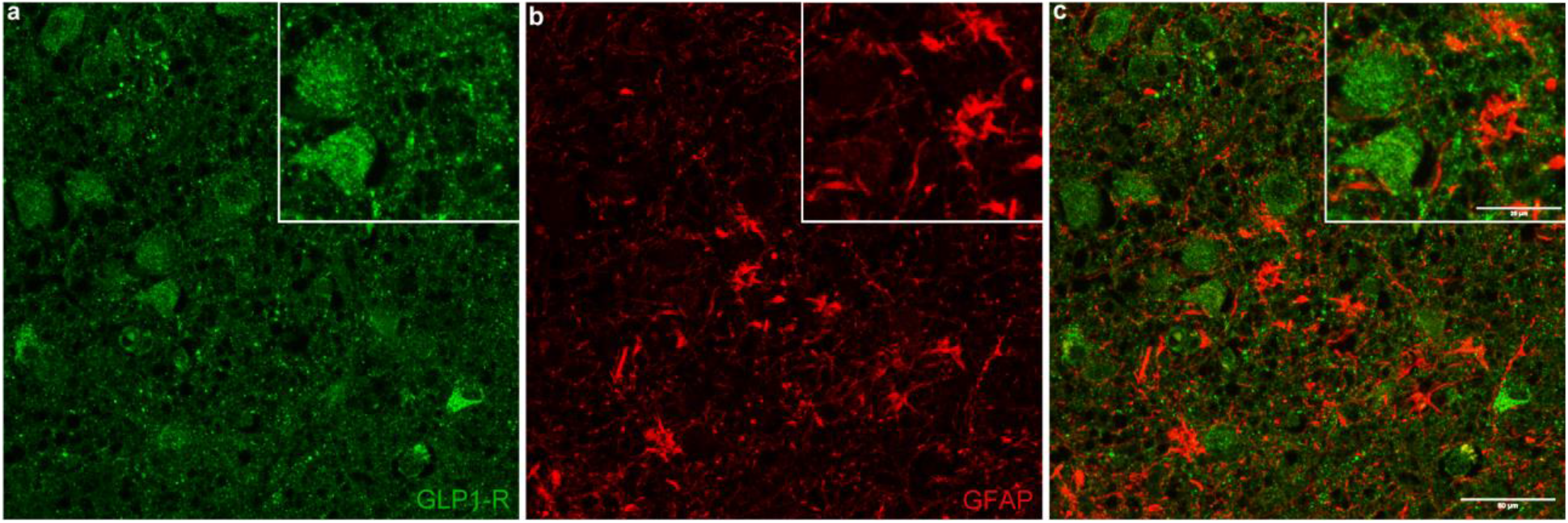
Confocal image showing GLP-1R and GFAP double immunofluorescence labeling in human hypothalamus. Panel (a) shows GLP-1R-immunoreactivity (green) and panel (b) displays GFAP-immunoreactivity (red). The merged image (c) shows the lack of colocalization of GLP-1R and GFAP. Scale bar equals 50 μm (25 μm in insets).

**Table 4:** Double immunofluorescence analysis of GLP-1-receptor-immunoreactive neurons colocalizing with NUCB2/nesfatin-1 in human hypothalamus. Numbers denote the percentage of GLP-1-receptor-immunoreactive neurons colocalized with NUCB2/nesfatin-1 (%).

|  | Case | BMI (Kg/m<sup>2</sup>) | PVN | SON | IFN | LH | BN |
| --- | --- | --- | --- | --- | --- | --- | --- |
| **Normal weight group** | 2 | 21.6 | 81.2 | 94.7 | 95.2 | 85.6 | 95.1 |
|  | 4 | 23.0 | 99.0 | 99.0 | n/a | 98.7 | 97.0 |
|  | 6 | 23.9 | 94.7 | 95.0 | 99.1 | n/a | n/a |
|  | 9 | 24.9 | 88.9 | 93.8 | 81.9 | 98.5 | 99.0 |
| **Mean value ± SEM** |  | **23.4 ± 0.7** | **91.0 ± 3.9** | **95.6 ± 1.2** | **92.1 ± 5.2** | **94.3 ± 4.3** | **97.0 ± 1.1** |
| **Non-normal weight group** | 14 | 29.0 | 94.5 | 93.8 | 85.5 | 99.0 | 90.4 |
|  | 18 | 29.4 | 99.9 | 99.4 | 99.0 | 99.5 | 99.0 |
|  | 23 | 31.7 | 85.2 | 98.9 | 95.5 | 98.7 | 91.7 |
|  | 25 | 32.4 | 86.0 | 99.0 | 71.4 | 99.0 | 99.0 |
| **Mean value ± SEM** |  | **30.6 ± 0.8** | **91.4 ± 3.5** | **97.8 ± 1.3** | **87.9 ± 6.2** | **99.0 ± 0.2** | **95.0 ± 2.3** |
| **Total Mean value ± SEM** |  | **27.0 ± 1.5** | **91.2 ± 2.4** | **96.7 ± 0.9** | **89.7 ± 3.9** | **97.0 ± 1.9** | **95.9 ± 1.4** |

Abbreviations: BMI, Body Mass Index; BN, basal nucleus; PVN, paraventricular nucleus; SON, supraoptic nucleus; IFN, infundibular nucleus; LH, lateral hypothalamic area; SEM, Standard Error of the Mean; n/a, not assessed.

**Table 5:** Double immunofluorescence analysis of NUCB2/nesfatin-1-immunoreactive neurons colocalizing with GLP-1 receptor in human hypothalamus. Numbers present the percentage of NUCB2/nesfatin-1-immunoreactive neurons colocalized with GLP-1 receptor (%).

|  | Case | BMI<br>(Kg/m<sup>2</sup>) | PVN | SON | IFN | LH | BN |
| --- | --- | --- | --- | --- | --- | --- | --- |
| **Normal weight group** | 2 | 21.6 | 60.6 | 72.2 | 61.0 | 63.6 | 72.9 |
|  | 4 | 23.0 | 62.9 | 46.6 | n/a | 70.3 | 77.5 |
|  | 6 | 23.9 | 55.2 | 68.4 | 82.0 | n/a | n/a |
|  | 9 | 24.9 | 58.1 | 42.3 | 60.1 | 64.7 | 71.8 |
| **Mean value ± SEM** |  | **23.4 ± 0.7** | **59.2 ± 1.7** | **57.4 ± 7.6** | **67.7 ± 7.2** | **66.2 ± 2.1** | **74.1 ± 1.7** |
| **Non-normal weight group** | 14 | 29.0 | 62.2 | 75.0 | 69.4 | 29.5 | n/a |
|  | 18 | 29.4 | 39.2 | 53.5 | 52.5 | 62.1 | 89.1 |
|  | 23 | 31.7 | 50.0 | 67.7 | 51.3 | 75.5 | 98.0 |
|  | 25 | 32.4 | 52.2 | 56.5 | 58.8 | 38.3 | 42.7 |
| **Mean value ± SEM** |  | **30.6 ± 0.8** | **50.9 ± 4.7** | **63.2 ± 5.0** | **58.0 ± 4.1** | **51.4 ± 10.6** | **76.6 ± 17.0** |
| **Total Mean value ± SEM** |  | **27.0 ± 1.5** | **55.1 ± 2.8** | **60.3 ± 4.3** | **62.2 ± 4.0** | **57.7 ± 6.5** | **75.3 ± 7.7** |

Abbreviations: BMI, Body Mass Index; BN, basal nucleus; PVN, paraventricular nucleus; SON, supraoptic nucleus; IFN, infundibular nucleus; LH, lateral hypothalamic area; SEM, Standard Error of the Mean; n/a, not assessed.

## Discussion

In the present study, we showed that GLP-1R protein is localized in several human hypothalamic nuclei implicated in the orchestration of energy metabolism and neuroendocrine and autonomic functions [26]. The most pronounced expression was observed in the PVN, SON, IFN, LH, and in cholinergic basal forebrain nuclei. GLP-1R-ir cells were also detected, although to a lesser extent, in other nuclei such as the VMH, DMH, TM, perifornical, medial preoptic, and mammillary as well as in bed nucleus of the stria terminalis. Importantly, in LH, GLP-1R protein expression was significantly lower in overweight or obese subjects compared with normal weight individuals and did negatively correlate with BMI. Interestingly, GLP-1R-immunopositive neurons extensively overlapped with the anorexigenic and anti-obesogenic neuropeptide NUCB2/nesfatin-1 in hypothalamic nuclei, namely PVN, SON, IFN, and LH, and in basal nucleus. In all the aforementioned nuclei, percentages of colocalization between GLP-1R and NUCB2/nesfatin-1 was not found to be altered in relation to BMI. Additionally, GLP-1R was not localized in hypothalamic GFAP-positive astrocytes.

GLP-1R localization, as described herein, is similar with previous studies in humans and animals [3–5, 10, 27–34]. According to our data, GLP-1R-ir cells, although in general agreement between studied cases, showed an interindividual variation in staining intensity in a few hypothalamic nuclei. These findings are consistent with previous reports in humans showing variations in hypothalamic GLP-1R mRNA and protein expression [4, 5]. The prominent immunoexpression of GLP-1R in the PVN, SON, IFN, and LH is in agreement with neuroanatomical evidence and functional analyses from human and animal studies, revealing that GLP-1R eating-inhibitory and anti-obesogenic effects are possibly mediated by various brain areas including hypothalamic nuclei [2–5, 10, 27–34]. In the present study, GLP-1R was also localized, although to a lesser extent, in perifornical nucleus, VMH, DMH, and TM, which have discrete roles in feeding behavior. GLP-1R immunoreactivity was also detected in the medial preoptic area, suprachiasmatic nucleus, mammillary nucleus, and bed nucleus of the stria terminalis, which participate in food intake modulation in response to temperature change, biological rhythms, memory, and emotional and behavioral responses to stress, respectively [26, 35–38]. The significance of this finding is unclear at present and further investigation is warranted. It can be hypothesized that GLP-1R might play a role in the behavioral act of feeding or in so far undefined physiological actions.

Furthermore, herein we showed that BMI had a significant moderate negative correlation with GLP-1R in the LH. Interestingly, GLP-1R immunoexpression was lower in overweight or obese subjects, compared with normal weight ones. As one of the most extensively interconnected brain areas, LH is involved in different aspects of food consumption, including hedonic feeding, and in body weight regulation [39, 40]. It should be noted that direct administration of GLP-1 into the LH of rats acutely suppresses feeding [41]. Additionally, the decrease in GLP-1R immunoreactivity in the LH of overweight or obese individuals shown herein, is comparable to the decreased expression of GLP-1R mRNA reported in the human PVN and IFN of type 2 diabetic patients [4]. It may be hypothesized that reduced GLP-1R in LH contribute to dysregulation of homeostatic and/or hedonic feeding behavior and consequently to body weight gain. Although it has been suggested that the effects of central GLP-1R activation on food intake and energy expenditure are mediated by at least partially separate hypothalamic areas and circuitries [2], it is unclear whether regional variations in LH GLP-1R expression, might influence GLP-1R signaling in hypothalamus. The factors contributing to these variations in hypothalamic nuclei and the concomitant functional implications in humans need to be further elucidated.

Interestingly, GLP-1R was also found to widely colocalize with the anorexigenic and anti-obesogenic neuropeptide NUCB2/nesfatin-1 in human hypothalamic nuclei involved in the regulation of energy metabolism, namely PVN, SON, IFN, and LH. In a previous study by Ten Kulve et al in the human hypothalamus. [4], GLP-1R was only sporadically colocalized with energy balance-related neuropeptides, such as NPY, AgRP, and POMC. It is intriguing to hypothesize that the extensive immunolocalization of GLP-1Rs on neurons expressing NUCB2/nesfatin-1, as observed herein, might indicate that GLP-1 affects energy homeostasis by acting on these hypothalamic neurons. Furthermore, it has been reported that nesfatin-1 hypothalamic neurons mediate GLP-1 anorexigenic effects in rodents [17, 18]. In addition, intraperitoneal administration of GLP-1 has been described to significantly increase activated nesfatin-1 neurons in hypothalamic nuclei, while GLP-1-induced food intake reduction has been significantly attenuated by pretreatment with intracerebroventricular administration of antisense nesfatin-1 [19]. As GLP-1 and NUCB2/nesfatin-1 share similar physiologic properties in energy homeostasis and body weight [2, 11, 15, 34], the aforementioned data, taken together with our observations, may be indicative of GLP-1 and nesfatin-1 possible interactions in the human hypothalamus.

Furthermore, we found no immunoexpression of GLP-1R in human hypothalamic astrocytes. GLP-1R localization in the human hypothalamus, by means of immunohistochemistry and *in situ* hybridization, has been previously reported only in neurons [4, 5]. On the other hand, GLP-1R expression has been described in a human astrocytic cell line [9]. In this latter study, astrocytes displayed GLP-1R after being subjected to mechanical or metabolic stress. However, the complexity of neuronal and non-neuronal compartments of brain circuits may more accurately be described in vivo and/or ex vivo compared to in vitro.

Interestingly, in our study, GLP-1R immunoexpression was observed in the cholinergic basal forebrain nuclei, a brain region known to participate in cognitive processes, memory, and feeding [26]. Although research has primarily focused on GLP-1R metabolic role, it is becoming increasingly obvious that GLP-1R activation has additional functions [2]. Experimental models and preclinical studies show beneficial effects of GLP-1R agonism on Alzheimer’s disease neuropathological features, including cholinergic basal forebrain nuclei degeneration, and on cognition [42–44]. Recent reports suggest that brain-gut peptides, including GLP-1 and nesfatin-1, share neuroprotective properties and might have therapeutic potential for neurodegenerative disorders [45–47]. The presence of GLP-1R in the human basal nucleus and its colocalization with nesfatin-1, as described in our study, is in line with this notion. However, the functional importance of these findings in the human cholinergic basal forebrain nuclei remains to be determined.

Animal models have strongly improved our understanding of hypothalamic responses as well as of factors contributing to body weight gain. It remains, however, difficult to assess whether these data can be extrapolated to neuroendocrine mechanisms underlying obesity in the human hypothalamus. Elucidating these neuronal pathways in humans entails technical difficulties and has limitations. Postmortem studies allow for a detailed anatomical and neurochemical examination of the energy balance-related circuitry, albeit causation cannot be determined. However, important biological information may be gleaned, providing an opportunity to therapeutically target obesity-associated maladaptive physiologic alterations in central neuronal networks. Further studies are warranted to clarify the functional significance of reduced GLP-1R protein expression in overweight or obese individuals in certain hypothalamic regions as well as any possible role of non-neuronal cells and the potential effects of nesfatin-1 in GLP-1R signaling in human hypothalamic circuits controlling energy homeostasis.

## Acknowledgments

The present study is dedicated to the memory of Professor of Anatomy-Histology-Embryology John Varakis who passed away on September 14, 2020, aged 79 years. Professor Varakis was an excellent mentor, colleague, and friend to students and scientific staff of University of Patras Medical School. We are sincerely grateful for his academic integrity and mental acuity which guided, inspired, and motivated us towards the search of scientific truth. The authors would also like to gratefully acknowledge the Advanced Light Microscopy Facility of Patras Medical School for sharing equipment.

## Data availability statement

The data that support the findings of this study are available from the corresponding author upon reasonable request.

---

**Supplementary Table 1:**
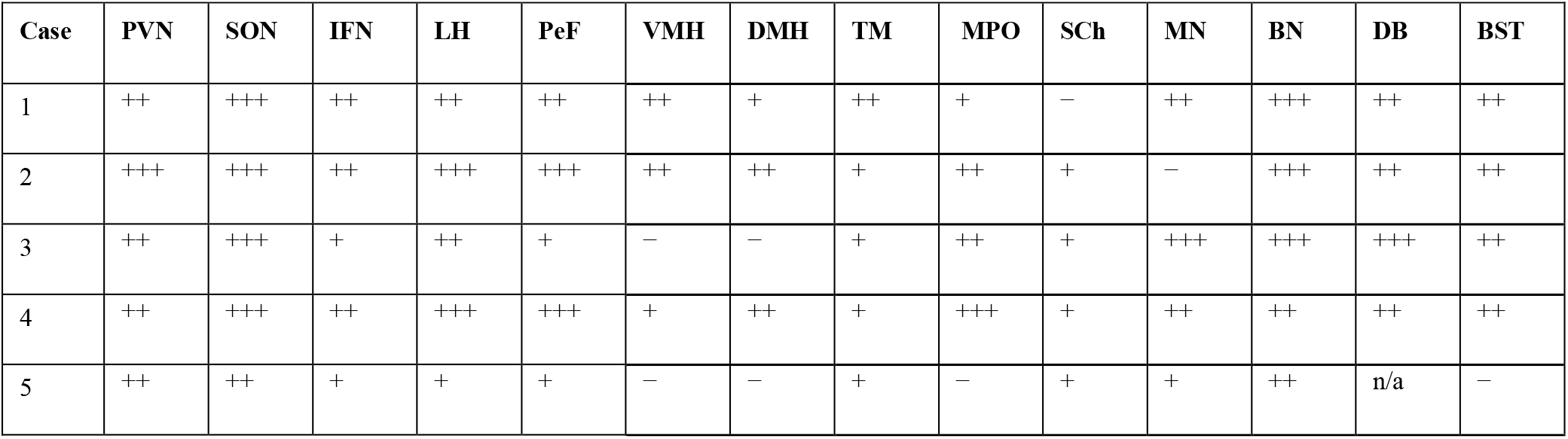
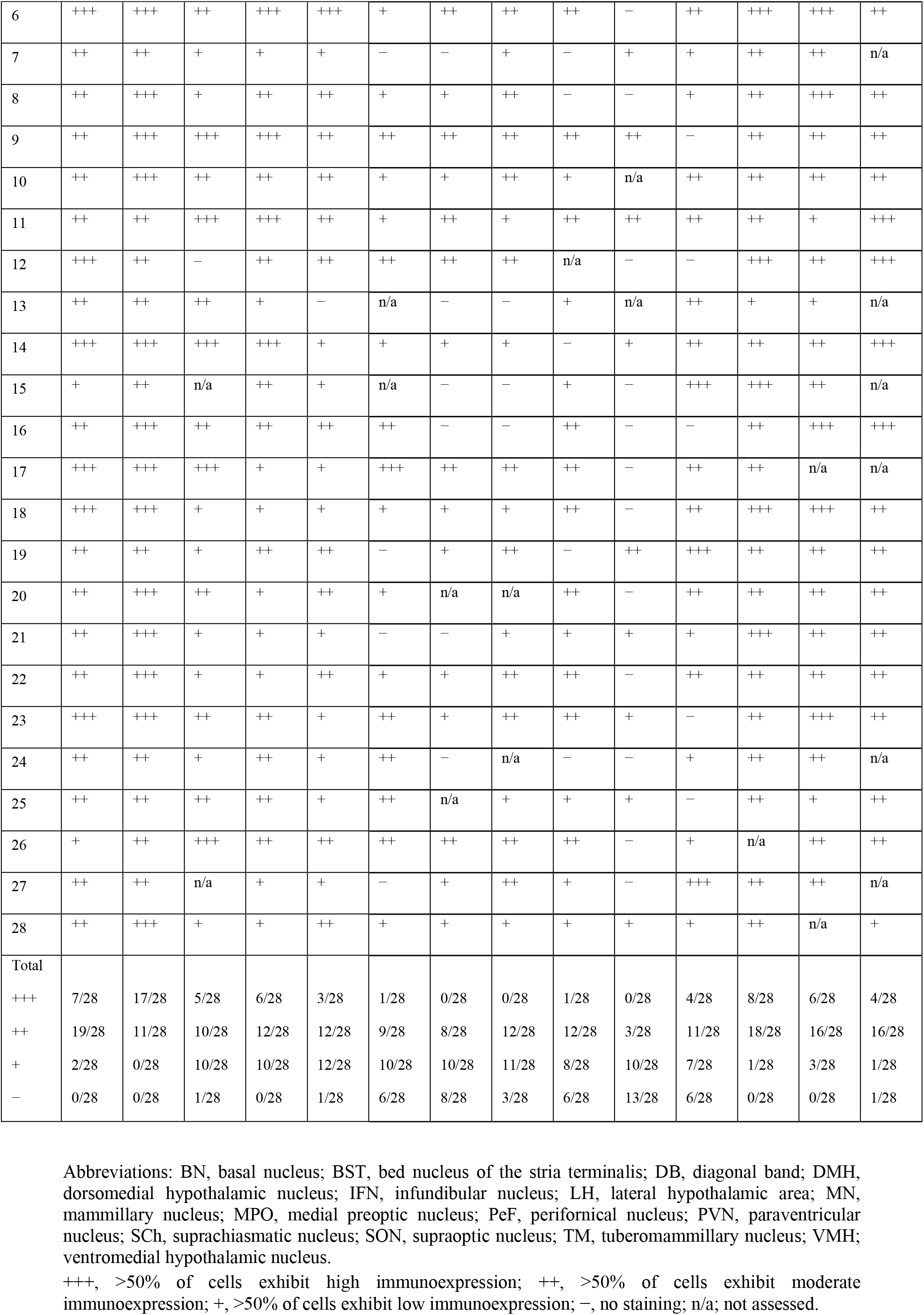
Distribution of GLP1R protein expression in the human hypothalamus of all examined cases, presented as immunostaining score, using rabbit polyclonal anti-GLP-1R antibody (Origene, Rockville, MD, USA, TA336864).

**Supplementary Table 2:**
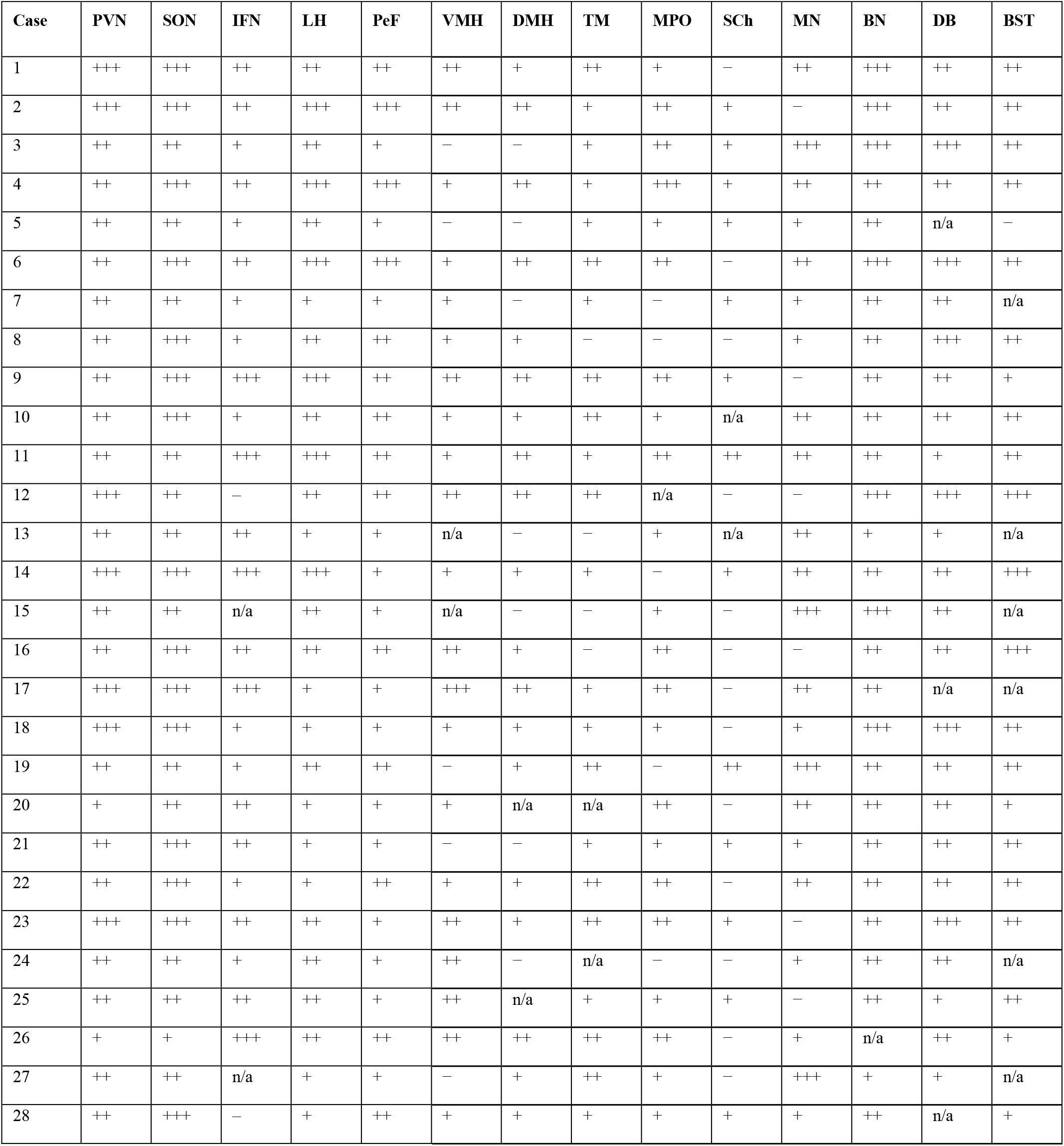
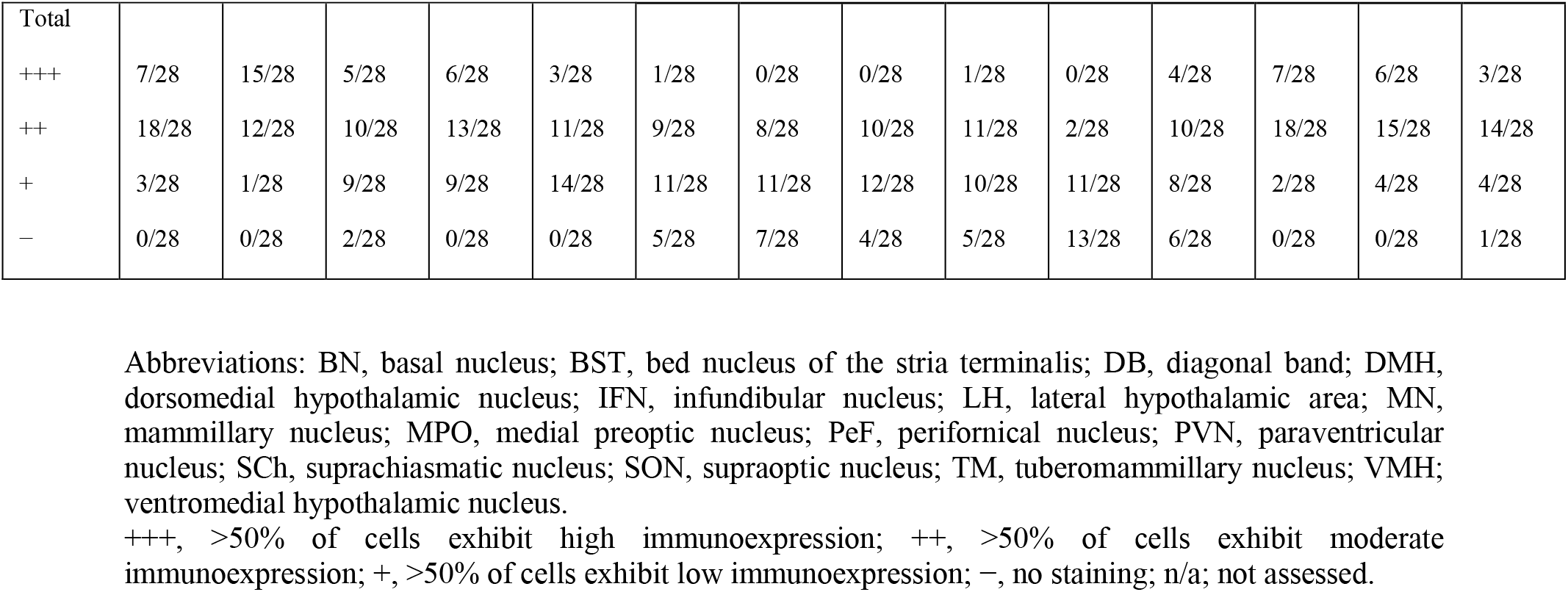
Distribution of GLP1R protein expression in the human hypothalamus of all examined cases, presented as immunostaining score, using rabbit polyclonal anti-GLP-1R antibody (Merck Millipore, Darmstadt, Germany, AB9433-I).

